# Fibroblast Protection of *Borrelia burgdorferi* from Doxycycline, Cefuroxime and Daptomycin Combination is Eliminated by Oregano or Carvacrol Essential Oil

**DOI:** 10.1101/861575

**Authors:** Chunxiang Bai, Hua Yang, Peng Cui, Rong Quan, Ying Zhang

## Abstract

*Borrelia burgdorferi* could be occasionally recovered from patients after antibiotic treatment, which indicates it may resist eradication by antibiotic and host defense mechanisms. Skin fibroblast cells have previously been shown to protect the killing of *B. burgdorferi* by ceftriaxone, a powerful antibiotic commonly used to treat Lyme disease. In this study, we evaluated if fibroblast cells could also protect against the doxycycline+ cefuroxime+ daptomycin drug combination which has previously been shown to completely eradicate highly persistent biofilm-like microcolonies of *B. burgdorferi.* To do so, we utilized a GFP-labeled *B. burgdorferi* for infection of murine fibroblast cells and assessed the effect of the drug combination on killing the bacteria in the presence or absence of the fibroblast cells. Surprisingly, we found that fibroblasts could protect *B. burgdorferi* from being completely killed by the drug combination doxycycline, cefuroxime and daptomycin, which eradicated *B. burgdorferi* completely in the absence of fibroblast cells. Interestingly, addition of essential oil carvacrol or oregano at 0.1% could enhance the activity of the doxycycline+ cefuroxime+ daptomycin drug combination and led to complete eradication of *B. burgdorferi* even in the presence of fibroblast cells. Further studies are needed to determine if the essential oil drug combinations could eradicate persistent *B. burgdorferi* infection in vivo in animal models. Our study provides a useful and convenient *ex vivo* model for evaluating different drug regimens needed for developing more effective treatment of persistent Lyme disease in the future.

## Introduction

Lyme disease is a multisystem infection caused by the spirochete *Borrelia burgdorferi (B. burgdorferi)* that is transmitted by Ixodes tick bites into the skin of animals and humans (1, 2). The skin of hosts is the key interface where the tick bites (3), and the epidermis with its keratinocytes represents the first barrier encountered by ticks. Ticks overcome this barrier using their biting mouthpieces to penetrate deeply into the skin and reach the dermis, where the saliva interacts with resident cells such as dermal dendritic cells, mast cells and fibroblasts. *B. burgdorferi* invades fibroblasts, and it is of interest to note that fibroblast cells have been shown to protect *B. burgdorferi* from killing by antibiotics such as ceftriaxone(4, 5).

Antibiotics, such as doxycycline, amoxicillin, or cefuroxime, are effective for the majority of Lyme disease cases (6–10). However, approximately 10–20% of the Lyme disease patients treated with antibiotics still experience persisting symptoms of fatigue, muscle aches, joint pain and neurologic impairment, that lasted for more than 6 months, a condition called Post-Treatment Lyme Disease Syndrome (PTLDS) (7). Viable *B. burgdorferi* has been isolated from the skin (11), cerebrospinal fluid (11, 12) and blood (13) of patients after antibiotic treatment for Lyme disease. In various animal models, such as mice, dogs and monkeys, it has been shown that the current Lyme antibiotic treatment with doxycycline or ceftriaxone are unable to completely eradicate the *B. burgdorferi* (14–16). In addition, in vitro studies have also demonstrated that *B. burgdorferi* could develop persister bacteria that are tolerant to or not effectively killed by the current Lyme antibiotics (17–19). Furthermore, *B. burgdorferi* develops variant forms such as round bodies and aggregated biofilm-like microcolonies which are persisters with increasing tolerance to antibiotics when the culture grows to late stationary phase or under stress conditions (20). These biofilm-like microcolony forms are not killed by the current Lyme antibiotics singly or in combination but can be eradicated by persister drug daptomycin in combination with doxycycline+cefuroxime (20).

In addition to chemical drugs such as antibiotics, natural products such as essential oils which exert broad antimicrobial activity against bacteria, fungi, viruses and parasites (21–25) could offer potential therapeutic options. The promising antimicrobial activity of essential oils could also be used with antibiotics to reduce drug side effects, toxicity, resistance to single agents and enhance the antibiotic activity of drugs against bacteria (21, 26). We have previously shown that some essential oils had stronger activity against stationary phase *B. burgdorferi* than the current antibiotics used for treating Lyme disease (27–29). Therefore, in this study, we evaluated 4 active essential oils (carvacrol, cinnamon bark, oregano and clove bud, cinnamaldehyde, allspice, hydacheim, myrrh, garlic and thyme white), which have been shown to have good activity against stationary phase *B. burgdorferi* (27), and their combinations with cefuroxime, doxycycline and daptomycin for their ability to kill *B. burgdorferi* in the fibroblast model.

## Materials and Methods

### Bacterial strain and culture condition

*B. burgdorferi* strain B31 expressing the green fluorescent protein (GFP) marker (GFP *B. burgdorferi)* was stored at −80°C, thawed at room temperature, and cultured at 33 °C in BSK-H medium. The cultures were monitored by microscopy for growth and contamination. The number of GFP-labeled *B. burgdorferi* was determined using a Petroff-Hausser bacteria counting chamber.

### Antibiotics and essential oils

Doxycycline (Dox), cefuroxime (CefU) (Sigma-Aldrich, USA) and daptomycin (Dap) (AK Scientific, Inc., USA) were dissolved in suitable solvents to form 5 mg/ml stock solutions. The antibiotic stocks were filter-sterilized by 0.2 μm filter and stored at −20°C. Essential oils (carvacrol, cinnamon bark, oregano and clove bud, cinnamaldehyde, allspice, hydacheim, myrrh, garlic and thyme white) were also dissolved in organic solvent dimethyl sulfoxide (DMSO) at 20%, followed by dilution to achieve desired dilutions as previously described (27).

### Cell cultures

Murine fibroblast NIH/3T3 cells (ATCC) were cultured in culture bottles with the area of 25 cm^2^ in Dulbecco’s modified Eagle medium (DMEM; GIBCO) containing 10% fetal bovine serum (FBS; GIBCO) at 37°C in a humidified incubator with 5% CO_2_. When the confluence of cells reached 70%-80%, confluent cell cultures were split with 0.25% trypsin-EDTA (GIBCO) and then 3T3 cells were seeded onto 24-well plates at 37°C in a humidified incubator with 5% CO_2_.

### Coculture of GFP-labeled *B. burgdorferi* and 3T3 cells

When the confluence stage was reached, GFP-labeled *B. burgdorferi* at stationary phase of growth (SP) was centrifuged at 10,000 rpm for 10 min and resuspended in DMEM supplemented with 10% FBS. Then GFP *B. burgdorferi* was added to tissue culture plate wells containing 3T3 cells in a 1:10 ratio of fibroblasts against the bacteria or to wells without cells (31). Cells and *B. burgdorferi* were cocultured for 48 h. Then the cells were then washed twice with warm PBS, fresh DMEM medium containing cefuroxime, doxycycline and daptomycin at 5 μg/mL or essential oils at three different concentrations 0.1%, 0.05% and 0.02% were added and the cocultures were further incubated for 7 days before the effect of the treatment was evaluated by microscopy or subculture study (5).

### Subculture study

After the cocultures were treated with antibiotics or essential oils for 7 days as described previously (5) the cells were washed once with warm PBS and lysed by adding 0.5 mL of distilled H_2_O for 5 min. Adherent cells were then scraped off, and the total content of each well was inoculated into 1 ml of BSK medium. Cultures were incubated at 33□ for 7 days or 21 days and monitored for the presence of viable *B. burgdorferi* by fluorescence microscopy. Control wells without cells but with spirochetes underwent the same procedure.

### Microscopy

The subcultured GFP *B. burgdorferi* was examined using BZ-X710 All-in-One fluorescence microscope (KEYENCE, Inc.). The residual cell viability reading was confirmed by using fluorescence microscopy. The toxic effect of essential oils on the fibroblast cells was evaluated after incubating the mixture for 7 days when the morphology of the fibroblast cells was observed using phase contrast function of the BZ-X710 All-in-One fluorescence microscope (KEYENCE, Inc.).

## Results

### Fibroblast cells protect *B. burgdorferi* from being killed by different antibiotics including drug combination Cefuroxime+ Doxycycline+ Daptomycin

Previous studies have shown that fibroblasts provided protection against ceftriaxone [5]. Here we evaluated whether a powerful persister drug combination CefU+ Dox + Dap, which is known to completely eradicate *B. burgdorferi* aggregated microcolony persisters in vitro (20) could eradicate *B. burgdorferi* from fibroblast cells. Surprisingly, as shown in Figure 1 and Table 1, the presence of 3T3 fibroblast cells protected the *B. burgdorferi* from being completely killed not only by single antibiotics doxycycline, cefuroxime, or daptomycin but also by the triple drug combination Cefuroxime+ Doxycycline+ Daptomycin as shown by residual green GFP labeled *B. burgdorferi* remaining after drug treatment for 7 days. However, we did find that Cefuroxime+ Doxycycline+ Daptomycin combination was more effective in killing *B. burgdorferi* than any other single antibiotics (Figure 1). Nevertheless, none of them including the triple drug combination Cefuroxime+ Doxycycline+ Daptomycin could completely kill *B. burgdorferi* as shown by visible spirochetal growth after 21-day subculture (Table 1).

**Figure 1.**
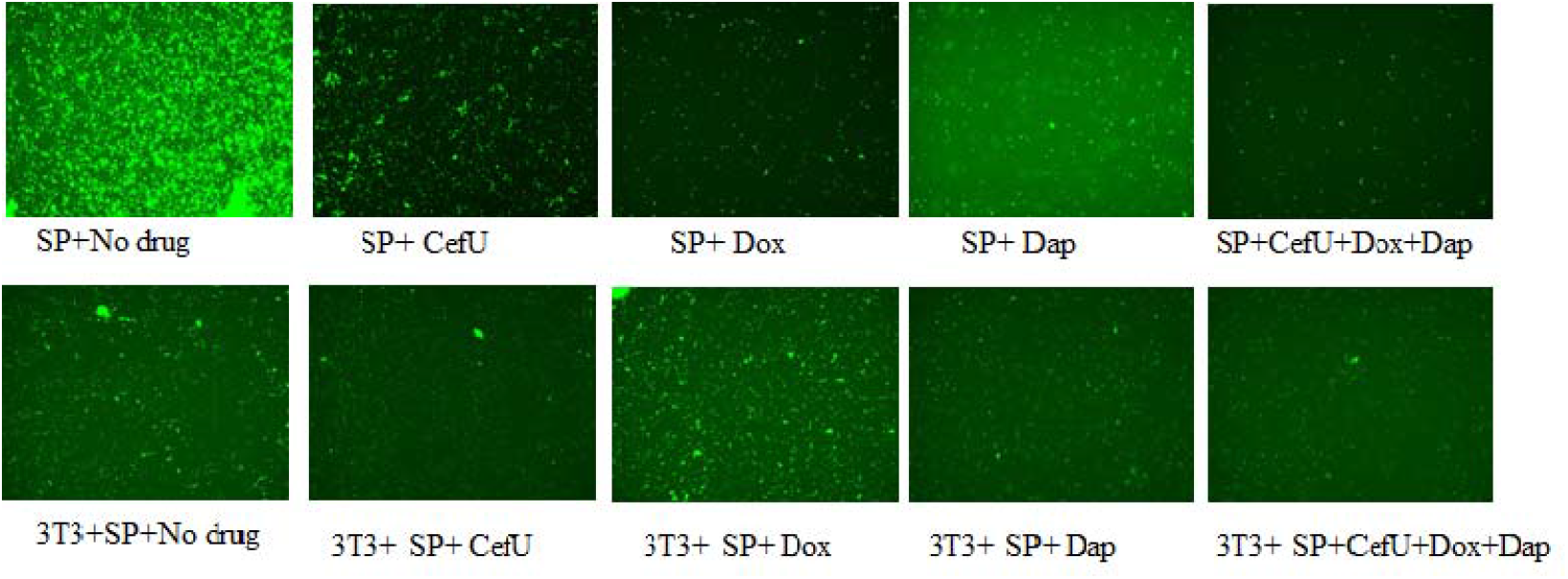
Coculture of GFP *B. burgdorferi* with 3T3 cells treated with antibiotics ex vivo. ^a^ Representative live images of stationary phase *B. burgdorferi* treated with different antibiotics at 200× magnification. *B. burgdorferi* at stationary phase of growth were added to wells containing a monolayer or to wells without cells (1.5×10^6^ live organisms/well) and incubated. Cells and *B. burgdorferi* were cocultured for 48 h. The cells were then washed twice with warm PBS, fresh tissue culture medium containing antibiotics at 5 μg/mL was added, and the cocultures were further incubated for 7 days. After 7 days, cultures were monitored for the presence of viable *B. burgdorferi.* Control wells without cells but with *B. burgdorferi* underwent the same procedure. Abbreviations: stationary phase culture of *B. burgdorferi* - SP, murine fibroblasts NIH/3T3 - 3T3, doxycycline - Dox, cefuroxime - CefU, daptomycin - Dap.

**Table 1.** Subculture the lysate of 3T3 infected with *B. burgdorferi* which were treated with antibiotics.

|  | Subculture for 21-days |
| --- | --- |
| SP+CefU | Growth |
| SP+Dox | Growth |
| SP+Dap | Growth |
| SP+CefU+Dox | Growth |
| SP+CefU+Dox+Dap | Growth |
| 3T3+SP+CefU | Growth |
| 3T3+SP+ Dox | Growth |
| 3T3+SP+Dap | Growth |
| 3T3+SP+CefU+Dox | Growth |
| 3T3+SP+CefU+Dox+Dap | Growth |

<sup>a</sup>After coculture of GFP *B. burgdorferi* with 3T3 cells treated with antibiotics *ex vivo* described as Figures 1, the total content of each well was inoculated into 1 ml of BSK-H medium. Cultures were incubated at 33°C for 21 days. After that, cultures were monitored for the presence of viable *B. burgdorferi* with microscope. Abbreviations: stationary phase culture of *B. burgdorferi* - SP, murine fibroblasts NIH/3T3 - 3T3, doxycycline - Dox, cefuroxime - CefU, daptomycin - Dap.

### Effect of essential oils on killing GFP *B. burgdorferi* in the fibroblast ex vivo model

We also evaluated essential oils for their activity to kill GFP *B. burgdorferi* cocultured with 3T3 fibroblast cells ex vivo. We tested 10 essential oils (carvacrol, cinnamon bark, oregano and clove bud, cinnamaldehyde, allspice, hydacheim, myrrh, garlic and thyme white), which were shown to have excellent activity against stationary phase *B. burgdorferi* in our previous studies (27, 28), at three different concentrations (0.1%, 0.05% and 0.02%) for activity against GFP *B. burgdorferi* cocultured with 3T3 cells. We found oregano oil and carvacrol (active component of oregano) at the lower concentration of 0.1% could kill all stationary phase *B. burgdorferi*, but cinnamon bark and clove bud could not eradicate all stationary phase *B. burgdorferi* (Figure 2A). At the same time, carvacrol and oregano at the concentration of 0.1% reduced significantly the number of stationary phase *B. burgdorferi* and showed stronger activity against *B. burgdorferi* in the presence of 3T3 cells than at the lower concentrations of 0.05% and 0.02%, but still could not completely eradicate all stationary phase GFP *B. burgdorferi* (Figure 2B). Meanwhile, oregano oil and carvacrol had no significant effect on cell morphology. Some essential oils such as myrrh and garlic had very serious toxicity effect on cell morphology (Table 2). Therefore, we combined oregano oil and carvacrol with the triple persister drug regimen (Dox+CefU+Dap) to evaluate whether the essential oil combinations could eradicate the persistent GFP-*B. burgdorferi* in the fibroblast cells.

**Figure 2.**
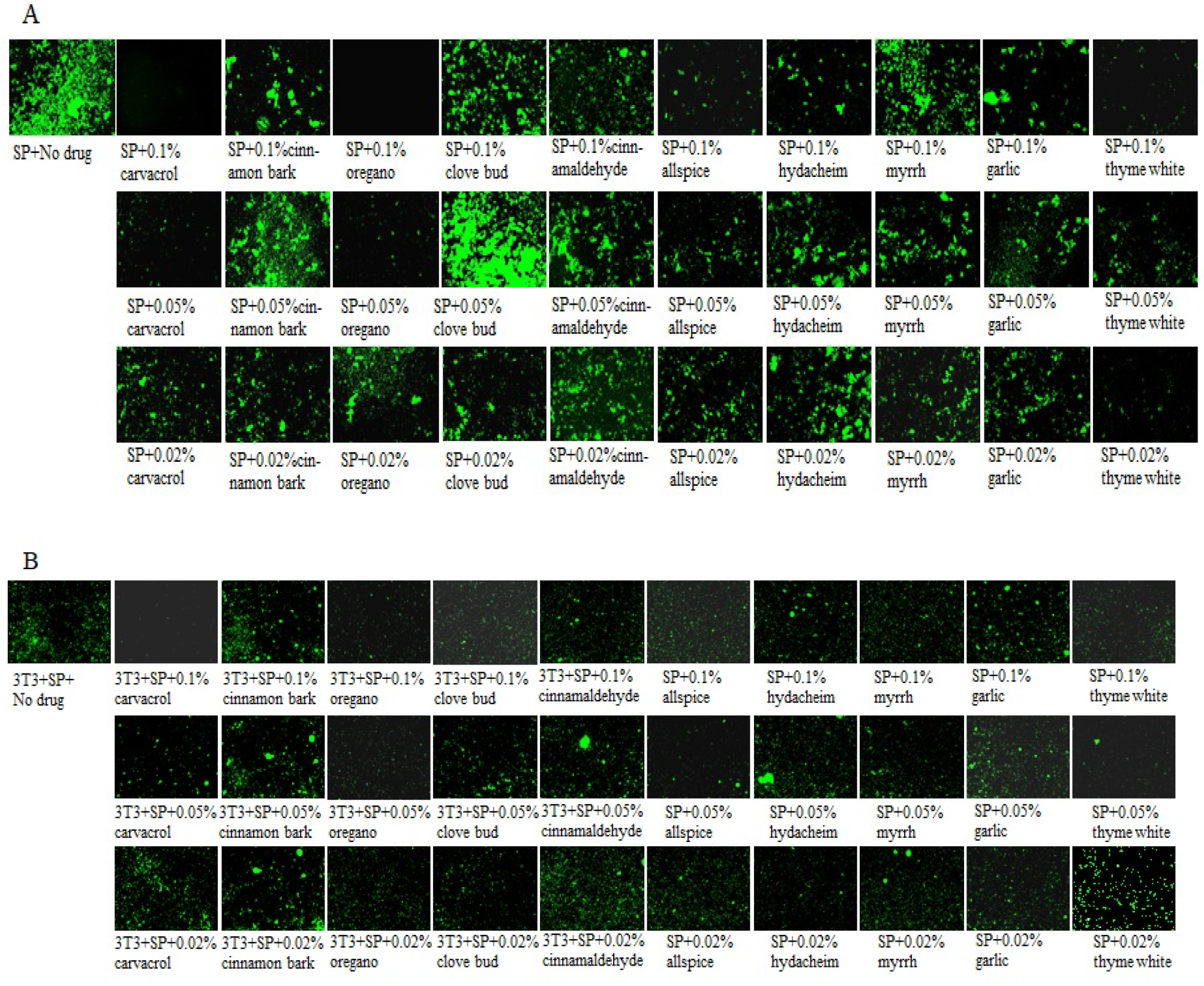
Coculture of GFP *B. burgdorferi* with 3T3 cells treated with essential oils ex vivo. Representative images of stationary phase *B. burgdorferi* treated with essential oils in the absence or presence of 3T3 fibroblast cells at 200 × magnification. We evaluated combination with essential oils ex vivo for activity against a 7-day-old *B. burgdorferi* stationary phase culture. *B. burgdorferi* at stationary phase of growth were added to wells containing a monolayer or to wells without cells (1.5×10^6^ live organisms/well) and incubated. Cells and *B. burgdorferi* were cocultured for 48 h. The cells were then washed twice with warm PBS, fresh tissue culture medium containing essential oils at a concentration of 0.1%, 0.05% and 0.02% were added the cocultures, and the cocultures were incubated for 7 days. After 7 days, cultures were monitored for the presence of viable *B. burgdorferi.* Control wells without cells but with *B. burgdorferi* underwent the same procedure. (A) Stationary phase culture of B. burgdorferi treated with essential oils. (B) Stationary phase culture of *B. burgdorferi* treated with essential oils. Abbreviations: stationary phase culture of *B. burgdorferi* - SP, murine fibroblasts NIH/3T3 - 3T3.

**Table 2.** Cytotoxicity effect of essential oils on 3T3 fibroblast cells.

| Essential oils |  | Cytotoxicity effect |
| --- | --- | --- |
| Carvacrol | 0.1% | No |
|  | 0.05% | No |
|  | 0.02 | No |
| Cinnamon bark | 0.1% | No |
|  | 0.05% | No |
|  | 0.02 | No |
| Oregano | 0.1% | No |
|  | 0.05% | No |
|  | 0.02 | No |
| Clove bud | 0.1% | No |
|  | 0.05% | No |
|  | 0.02 | No |
| Cinnamaldehyde | 0.1% | No |
|  | 0.05% | No |
|  | 0.02 | No |
| Allspice | 0.1% | No |
|  | 0.05% | No |
|  | 0.02 | No |
| Hydacheim | 0.1% | Yes |
|  | 0.05% | No |
|  | 0.02 | No |
| Myrrh | 0.1% | Yes |
|  | 0.05% | Yes |
|  | 0.02 | No |
| Garlic | 0.1% | Yes |
|  | 0.05% | Yes |
|  | 0.02 | No |
| Thyme white | 0.1% | No |
|  | 0.05% | No |
|  | 0.02 | No |

<sup>a</sup>3T3 cells were incubated with essential oils (0.1%, 0.05% and 0.02%) for 7 days to evaluate their potential toxicity on 3T3 cells. Cell morphology was observed by phase contrast microscopy.

### Essential oils in combination with the triple persister antibiotic regimen (Dox+CefU+Dap) completely eradicate GFP *B. burgdorferi* in the fibroblast cell model

To assess if essential oils could potentially enhance the activity of the persister drug combination, we selected oregano oil and carvacrol at the concentration of 0.1% and combined them with CefU, Dox and Dap to assess their activity on GFP *B. burgdorferi* in coculture with 3T3 cells. The results showed that while carvacrol, oregano or CefU+ Dox + Dap alone had good activity against *B. burgdorferi* in 3T3 cells, none of them alone was able to completely eradicate *B. burgdorferi* in the fibroblast cell model. Importantly, carvacrol or oregano in combination with CefU, Dox and Dap could kill all *B. burgdorferi* cells in the absence and even in the presence of fibroblast cells (Figure 3). In the absence of fibroblast cells, there was no *B. burgdorferi* after treatment with carvacrol or oregano (Figure 3). Subculture study also confirmed the high activity of carvacrol or oregano at 0.1% combined with CefU, Dox and Dap for complete eradication of GFP *B. burgdorferi* in the fibroblast cell model as evidence by no regrowth in 21 days. However, regrowth occurred in samples treated with carvacrol, oregano or CefU+ Dox + Dap alone in the fibroblast cell model. In contrast, in the absence of the fibroblast cells, no regrowth was observed in the samples treated with carvacrol or oregano or CefU, Dox and Dap after 21-day subculture (Table 3).

**Figure 3.**
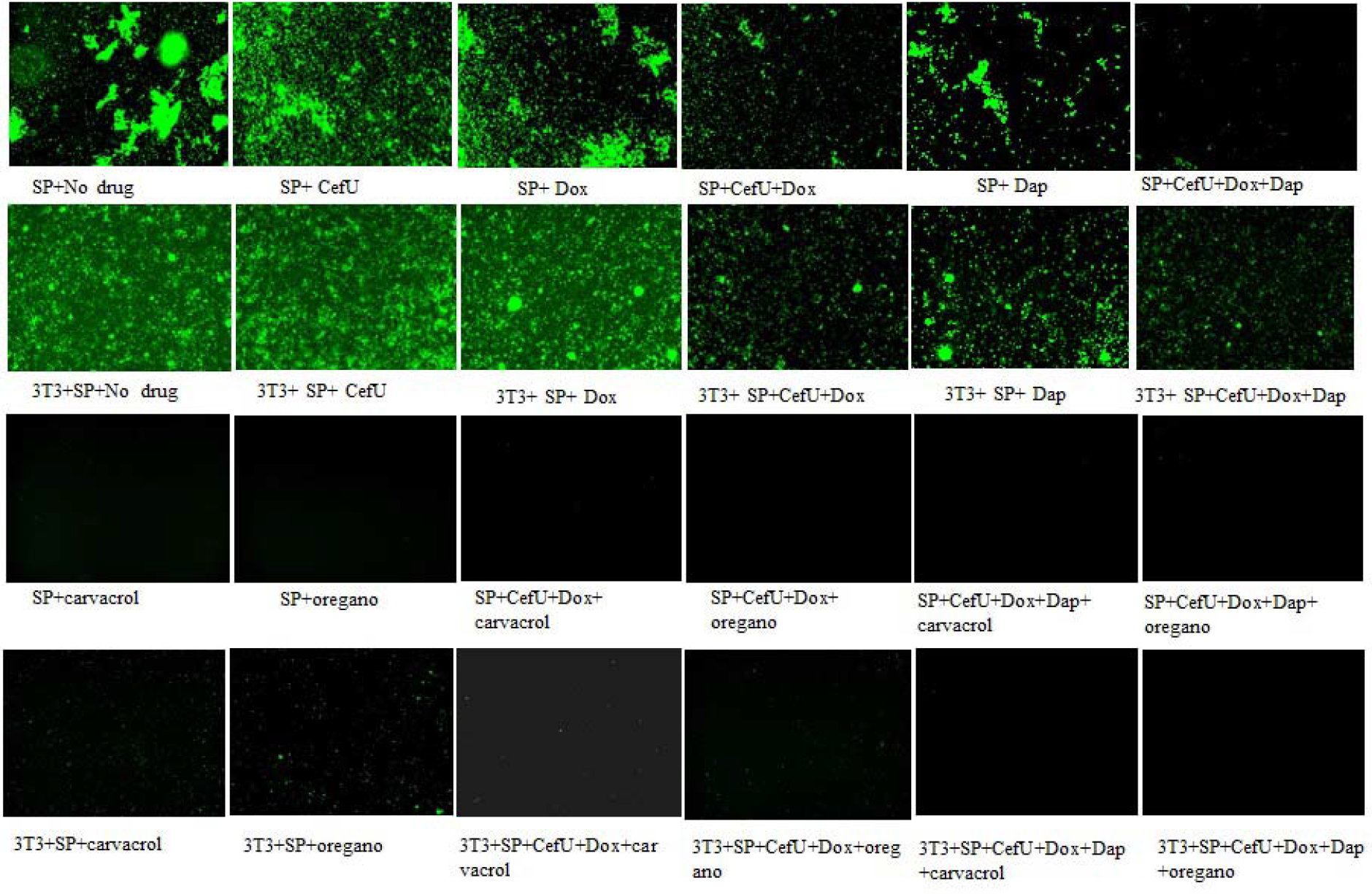
Coculture of GFP *B. burgdorferi* with 3T3 cells treated with antibiotics and essential oils ex vivo. ^a^Representative images of stationary phase *B. burgdorferi* treated with antibiotics and essential oils ex vivo at 200× magnification. We evaluated combination of antibiotics and essential oils ex vivo for activity against a 7-day-old *B. burgdorferi* stationary phase culture. Stationary phase *B. burgdorferi* cells were added to wells containing a monolayer of fibroblast cells (1.5×10^6^ organisms/well) or to wells without cells and incubated at 37°C. Cells and *B. burgdorferi* were cocultured for 48 h. The cells were then washed twice with warm PBS, fresh tissue culture medium containing antibiotics at 5 μg/mL or essential oils at a concentration of 0.1% were added, and the cocultures were incubated for 7 days. After 7 days, cultures were monitored for the presence of viable *B. burgdorferi.* Control wells without cells but with *B. burgdorferi* underwent the same procedure. Abbreviations: stationary phase culture of *B. burgdorferi* - SP, murine fibroblasts NIH/3T3 - 3T3, doxycycline - Dox, cefuroxime - CefU, daptomycin - Dap.

**Table 3.** Subculture the lysate of 3T3 infected with *B. burgdorferi* which were treated with antibiotics and essential oils.

|  | Subculture for 21-days |
| --- | --- |
| SP+CefU | Growth |
| SP+Dox | Growth |
| SP+Dap | Growth |
| SP+CefU+Dox | Growth |
| SP+CefU+Dox+Dap | Growth |
| SP+Carvacrol | No growth |
| SP+ Oregano | No growth |
| SP+CefU+Dox+ Carvacrol | No growth |
| 3T3+SP+CefU+Dox+ Oregano | No growth |
| SP+CefU+Dox+Dap+Carvacrol | No growth |
| SP+CefU+Dox+Dap+Oregano | No growth |
| 3T3+SP+CefU | Growth |
| 3T3+SP+ Dox | Growth |
| 3T3+SP+Dap | Growth |
| 3T3+SP+CefU+Dox | Growth |
| 3T3+SP+CefU+Dox+Dap | Growth |
| 3T3+SP+ Carvacrol | Growth |
| 3T3+SP+ oregano | Growth |
| 3T3+SP+CefU+Dox+ Carvacrol | Growth |
| 3T3+SP+CefU+Dox+ Oregano | Growth |
| 3T3+SP+CefU+Dox+Dap+ Carvacrol | No growth |
| 3T3+SP+CefU+Dox+Dap+ Oregano | No growth |

<sup>a</sup>After coculture of GFP *B. burgdorferi* with 3T3 fibroblast cells treated with antibiotics and essential oils *ex vivo*, the total content of each well was inoculated into 1 ml of BSK-H medium to determine if any residual bacteria not killed would regrow. Cultures were incubated at 33°C for 21 days and monitored for the presence of viable *B. burgdorferi* with fluorescence microscope. Abbreviations: SP- stationary phase culture of *B. burgdorferi*, 3T3- NIH/3T3 murine fibroblast cell line, Dox- doxycycline, CefU- cefuroxime, Dap- daptomycin.

## Discussion

In this study, we used stationary phase *B. burgdorferi*- fibroblast coculture model ex vivo to evaluate whether fibroblasts would protect *B. burgdorferi* from the action of some drugs and whether combinations of drugs and essential oils could eradicate *B. burgdorferi.* The reason why we chose this model are as follows. Firstly, in tuberculosis treatment, persister drug pyrazinamide is more active against stationary phase cells and persisters than against log phase growing cells and could shorten the therapy (32), which justifies the use of stationary phase *B. burgdorferi* enriched in persisters as a persister model. Secondly, *B. burgdorferi* is transmitted firstly into the skin by ticks, where it spreads locally and establishes infection (33, 34). Fibroblasts reside in the skin and interact with *B. burgdorferi.* Thirdly, fibroblasts have been shown to protect *B. burgdorferi* from the action of antibiotics commonly used for its eradication, especially the most powerful Lyme antibiotic ceftriaxone (5). This implies that fibroblast interaction with *B. burgdorferi* may allow persisters to develop that are not killed easily by the current Lyme antibiotics.

In this study, we were interested to know whether fibroblasts could also protect *B. burgdorferi* from the action of the powerful persister drug combination CefU, Dox and Dap, which was previously shown to completely eradicate *B. burgdorferi* persisters not killed by the current Lyme antibiotics in vitro (20). Surprisingly, the presence of the fibroblast cells indeed protected the *B. burgdorferi*, from eradication by the persister drug combination CefU, Dox and Dap as shown by regrowth in subculture study. Nevertheless, we did find that CefU, Dox and Dap combination was better against *B. burgdorferi* than each single antibiotics in the fibroblast cell persistence model, since none of them could completely kill *B. burgdorferi* as they were able to regrow and were still visible after 21-day subculture. We then tested 4 active essential oils carvacrol, cinnamon bark, oregano and clove bud which previously were shown to be highly active against *B. burgdorferi* (27) in the *B. burgdorferi-fibroblast* cell coculture model. We found that fibroblasts protected *B. burgdorferi* from eradication by the triple drug combination or carvacrol or oregano at 0.1% as either alone was unable to completely clear the bacteria. Interestingly, we found that when the triple drug combination doxycycline+cefuroxime+daptomycin was combined with essential oils carvacrol or oregano at 0.1%, all *B. burgdorferi* cells were completely eradicated such that the cells did not grow back in 21-day subculture. This indicates the potential value of essential oils when used in combination with antibiotics. Further studies are needed to confirm this in animal studies.

It has been previously demonstrated that *B. burgdorferi* penetrated endothelial cells inside and between these cells (35), or penetrated cultured human umbilical vein endothelial cells (36), but the viability of the intracellularly located spirochetes was not assessed. In our study, we were mainly focused on evaluating the effect of drug combination and essential oils on eradicating *B. burgdorferi* in the fibroblast persistence model, and whether the GFP *B. burgdorferi* is located inside or on the surface of the fibroblast cells is less clear. Future studies are required to precisely define the locale of the protective effect of fibroblasts for *B. burgdorferi* as well as whether other drugs with different modes of action and different ability to penetrate cells would eliminate the spirochetes even in the presence of fibroblast cells. One possibility that may explain why fibroblasts protected *B. burgdorferi* from the drug combination is that association of *B. burgdorferi* with fibroblast cells allows the bacteria to change the metabolic status and develop more persisters. Alternatively, association of *B. burgdorferi* with fibroblast allows the bacteria to attach to the fibroblast and facilitate biofilm formation such that even the powerful triple drug combination is still unable to eradicate the bacteria. It is also possible that *B. burgdorferi* could penetrate cells and survive in fibroblasts as shown by previous studies (36) and the intracellular survival would provide protection against antibiotics, since many antibiotics are much less concentrated in the cells than in extracellular spaces. In contrast, essential oils carvacrol and oregano due to their lipophilicity may have a better cell membrane association and penetration as well as intracellular distribution, which may explain the more effective killing and eradication of *B. burgdorferi* by essential oils carvacrol or oregano combined with the triple drug combination doxycycline, cefuroxime, daptomycin even in the presence of the fibroblast cells. Further studies are needed to elucidate the mechanisms involved.

In summary, the results in this study showed host fibroblast cells could protect *B. burgdorferi* from the killing by drug combination doxycycline, cefuroxime and daptomycin, but addition of essential oil carvacrol or oregano at 0.1% to this drug combination could completely eradicate *B. burgdorferi* with no regrowth. Future studies will be carried out to assess their safety and efficacy against *B. burgdorferi* infection in animal models.

## Acknowledgments

We thank Jon Skare for providing the GFP-labeled *B. burgdorferi.* This study was supported in part by grants from Department of Defense (W81XWH-17-1-0664), Global Lyme Alliance, and Steven & Alexandra Cohen Foundation.

## Conflicts of Interest

The authors declare no conflict of interest.

